# *minimap2-fpga*: Integrating hardware-accelerated chaining for efficient end-to-end long-read sequence mapping

**DOI:** 10.1101/2023.05.30.542681

**Authors:** Kisaru Liyanage, Hiruna Samarakoon, Sri Parameswaran, Hasindu Gamaarachchi

**Affiliations:** School of Computer Science and Engineering, University of New South Wales, Sydney, NSW, Australia; Genomics Pillar, Garvan Institute of Medical Research, Sydney, NSW, Australia; Centre for Population Genomics, Garvan Institute of Medical Research and Murdoch Children’s Research Institute, Australia; School of Electrical and Information Engineering, University of Sydney, Sydney, NSW, Australia

## Abstract

*minimap2* is the gold-standard software for reference-based sequence mapping in third-generation long-read sequencing. While *minimap2* is relatively fast, further speedup is desirable, especially when processing a multitude of large datasets. In this work, we present *minimap2-fpga*, a hardware-accelerated version of *minimap2* that speeds up the mapping process by integrating an FPGA kernel optimised for chaining. We demonstrate speed-ups in end-to-end run-time for data from both Oxford Nanopore Technologies (ONT) and Pacific Biosciences (PacBio). *minimap2-fpga* is up to 79% and 53% faster than *minimap2* for ∼ 30× ONT and ∼ 50× PacBio datasets respectively, when mapping without base-level alignment. When mapping with base-level alignment, *minimap2-fpga* is up to 62% and 10% faster than *minimap2* for ∼ 30× ONT and ∼ 50× PacBio datasets respectively. The accuracy is near-identical to that of original *minimap2* for both ONT and PacBio data, when mapping both with and without base-level alignment. *minimap2-fpga* is supported on Intel FPGA-based systems (evaluations performed on an on-premise system) and Xilinx FPGA-based systems (evaluations performed on a cloud system). We also provide a well-documented library for the FPGA-accelerated chaining kernel to be used by future researchers developing sequence alignment software with limited hardware background.

## Introduction

Third-generation long-read sequencing has gained significant popularity in the last few years and was recently acclaimed as the Method of the Year 2022^1^. In numerous long-read sequence analysis pipelines, *minimap2*^2^ is used as the gold standard tool for sequence mapping/alignment. Notably, *minimap2*^2^ is the preferred choice for sequence mapping in pipelines offered by leading third-generation sequencing companies, such as ONT (Guppy aligner) and PacBio (pbmm2). Due to *minimap2*’s ubiquitous use, several previous studies have attempted to accelerate *minimap2*^3–6^. However, most of these efforts have focused on optimising isolated parts of *minimap2* in a quarantined environment (not end-to-end integrated), with the only exception being CPU-based optimisations (using Intel AVX-512 instructions) by Kalikar et al.^3^. In this paper, we present *minimap2-fpga*, a Field Programmable Gate Array (FPGA) based hardware-accelerated version of *minimap2* that is end-to-end integrated. *minimap2-fpga* speeds up the mapping process of the widely used and routinely performed bioinformatics workflow - reference-based sequence mapping. *minimap2-fpga* supports mapping reads from both ONT and PacBio (though giving a higher speed-up for ONT read mapping).

The acceleration of the chaining step within *minimap2* has been explored in numerous previous studies, given its status as a key computational bottleneck within the software^3–6^. Sadasivan et al.^4^ used GPUs for accelerating this chaining step. Guo et al.^5^ accelerated the chaining step using FPGA, mainly targeting the read overlap mapping functionality in *minimap2*. Our previous work^6^ accelerated chaining using FPGAs targeting human reference-based mapping functionality in *minimap2*. These three works that use loosely coupled accelerator platforms (GPUs, FPGAs), only accelerate chaining in isolation and end-to-end integration is not performed with *minimap2*, thus limiting their practical utility. In contrast, Kalikar et al.^3^ use tightly coupled AVX-512 vector extensions available in modern Intel CPUs to accelerate chaining and integrate the acceleration to *minimap2*.

The isolated chaining step accelerator in our previous work^6^ combined a pipelined and parallel FPGA-based hardware architecture with a multi-threaded software execution of the chaining step computation. The system was able to split chaining tasks between CPU (multi-threaded) and FPGA (multi-kernel) platforms efficiently and process them in parallel to speed up the computation. Moreover, we demonstrated its capability to effectively process large-scale and realistic datasets frequently encountered in bioinformatics workflows. In this work, we integrate our previous FPGA-based hardware architecture^6^ into *minimap2* to enable end-to-end program execution, while exposing the accelerated chaining kernel as a well-documented software library.

Integrating the FPGA kernel into *minimap2* posed significant challenges when compared to designing the FPGA core presented in our prior work^6^. While data transfer overhead was previously considered^6^, the integration process exposed additional scheduling overheads when working with the multi-threaded execution environment in *minimap2*, resulting in marginal end-to-end speedup. In this paper, we present optimisation techniques for the splitting of chaining computations between the hardware accelerator and the CPU, while efficiently scheduling hardware-chosen computations on the accelerator, to achieve further acceleration.

We also discuss an unexpected caveat that arose when integrating the design, impacting final mapping accuracy, a factor not apparent in isolated testing environments. We detail how the FPGA design was modified to mitigate this issue. We believe that our findings will prove valuable for other bioinformatics tools requiring heterogeneous computing accelerations, emphasising the importance of integrating accelerated kernels back into software workflows. Our work highlights the potential for disruptive challenges which arise when integrating isolated acceleration of a single algorithm in a complex software environment, underpinning the need for end-to-end system integration.

Our end-to-end integrated *minimap2-fpga* requires a heterogeneous computing system that contains a CPU connected to an FPGA through PCI-Express. FPGA from the mainstream FPGA vendors Xilinx (now under AMD) and Intel (former Altera) are supported. Further, we have ensured that our implementation works on both locally set up (on-premises) computing systems and the systems available on the cloud (Amazon AWS and Intel DevCloud). *minimap2-fpga* is available at https://github.com/kisarur/minimap2-fpga and the accelerated chaining library called *chain-fpga* is available at https://github.com/kisarur/chain-fpga.

## Results

### minimap2-fpga and chain-fpga

*minimap2-fpga* is an FPGA-accelerated version of the *minimap2* software designed for reference-based sequence mapping. *minimap2-fpga* can be run using the same commands as the original *minimap2* software. The repository has separate branches for different FPGA platforms. The *xilinx* branch of the repository contains the source code for Xilinx FPGA, while the *intel* branch contains the source code for Intel FPGA. Despite being developed using platform-agnostic OpenCL, the build workflows and HLS pragmas used to generate efficient hardware differed between the two platforms, resulting in the need for separate branches. To enhance user convenience, we provide binaries for two platforms: 1. Intel Arria 10GX FPGA and 2. Xilinx UltraScale+ VU9P based FPGA available on cloud AWS EC2 F1 instances. An example of how a user can set up and execute *minimap2-fpga* on an AWS cloud (*f1*.*2xlarge* instance loaded with *FPGA Developer AMI Version 1*.*12*.*1*) is as below:

~~~
# setup the environment
git clone https://github.com/aws/aws-fpga.git $AWS_FPGA_REPO_DIR source $AWS_FPGA_REPO_DIR/vitis_runtime_setup.sh
# download github repo and build host application
git clone https://github.com/kisarur/minimap2-fpga.git
cd minimap2-fpga
make host
# run minimap2-fpga
./minimap2-fpga [minimap2 arguments]
~~~

*chain-fpga* is the isolated library API for the FPGA-based chaining which could be used in various bioinformatics tool development where chaining is a critical computational hotspot (e.g., sigmap^7^). This library API offers an easy-to-use interface for integrating the chaining accelerator into other bioinformatics tools without requiring expertise in custom hardware development. Similar to the *minimap2-fpga* repository, this repository also contains two separate branches for Xilinx and Intel FPGA, along with pre-compiled binaries. The following are the three key API functions supported by the library API:

~~~
// to initialize the OpenCL hardware resources
hardware_init()
// to perform core chaining on the FPGA-based hardware accelerator perform_core_chaining_on_fpga(args)
// to clean up the hardware resources allocated cleanup()
~~~

### Performance Comparison

The end-to-end run time of *minimap2-fpga* was benchmarked on a cloud Xilinx FPGA system and an on-premises Intel FPGA system. To represent realistic workloads, two large and complete datasets were employed for the benchmarks: a ∼ 30× ONT human dataset and a ∼ 50× PacBio CCS human dataset (see Materials and Methods). This intentional selection distinguishes our work from previous hardware-accelerated systems, which only performed benchmarks on small datasets, possibly due to challenges in managing memory effectively. We compared the run time of *minimap2-fpga* with the original *minimap2* software in two different modes for each system and dataset: one with base-level alignment enabled (w/ alignment) and the other without (w/o alignment).

For nanopore data (Figure 1a), *minimap2-fpga* is 79% faster than *minimap2* on the on-premise Intel FPGA system and 72% faster than *minimap2* on the cloud Xilinx FPGA system when mapping without base-level alignment. When mapping with base-level alignment enabled, *minimap2-fpga* shows a speedup of 62% and 40% on the on-premise and cloud systems, respectively. On PacBio data (Figure 1b), *minimap2-fpga* is 53% faster than *minimap2* on the on-premise Intel FPGA-based system and 31% faster on the cloud Xilinx FPGA-based system when mapping without base-level alignment. With base-level alignment enabled, *minimap2-fpga* shows a speedup of 10% and 7% on the on-premise Intel and cloud Xilinx systems, respectively.

**Figure 1.**
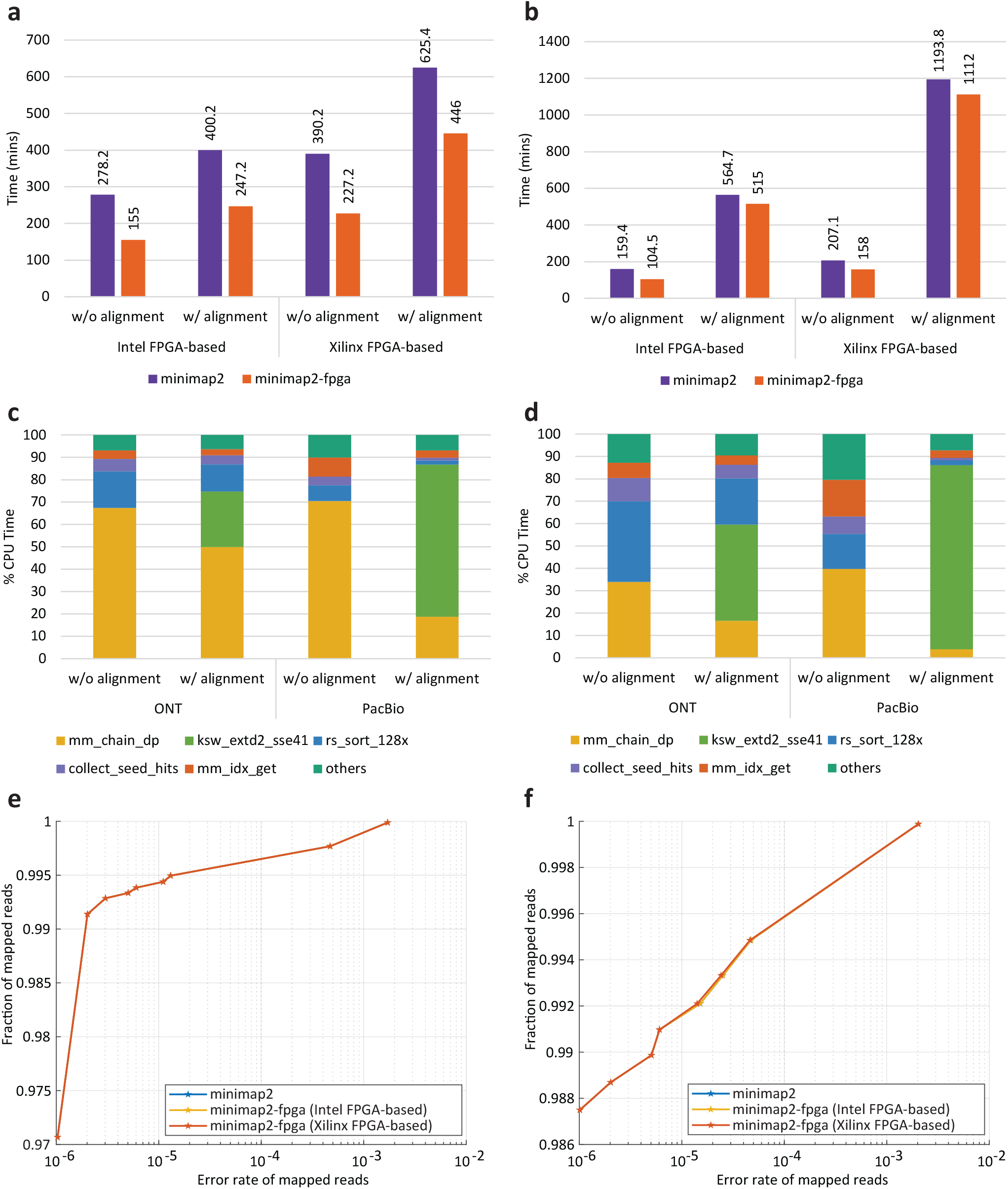
Run-time performance, function-level profiling and accuracy comparisons of *minimap2* vs. *minimap2-fpga*. (**a**) Run-time performance results of original *minimap2* vs. *minimap2-fpga* for ONT ∼ 30× dataset. (**b**) Run-time performance results of original *minimap2* vs. *minimap2-fpga* for PacBio ∼ 50× dataset. (**c**) Function-level profiling results of original *minimap2*. Profiling was done with the Perf profiler^8^ on Intel FPGA-based system for subsets of ONT and PacBio datasets. (**d**) Function-level profiling results of *minimap2-fpga*. Profiling was done with the Perf profiler^8^ on Intel FPGA-based system for subsets of ONT and PacBio datasets. (**e**) Accuracy comparison performed with simulated ONT reads for original *minimap2* vs. *minimap2-fpga* with base-level alignment. (**f**) Accuracy comparison performed with simulated ONT reads for original *minimap2* vs. *minimap2-fpga* without base-level alignment.

The variation in performance improvement across different datasets and modes can be attributed to the proportion of time spent on chaining (i.e. “mm_chain_dp” function which is the FPGA-accelerated component), out of the total execution time (Figure 1c). For nanopore data, original *minimap2* spends 67% of its time on chaining when performed without base-level alignment (Figure 1c), and this is reduced to 34% after FPGA acceleration (Figure 1d). When nanopore data is processed with base-level alignment, the chaining component on original *minimap2* takes up 50% of the total time, but this drops to 17% after being accelerated on FPGA. For PacBio data, the original *minimap2* spends 71% of its total execution time on chaining when mapping without base-level alignment, which is reduced to 40% after FPGA acceleration. When PacBio data is processed with base-level alignment, *minimap2* spends only 19% of its total execution time on chaining, which is reduced to 4% after being accelerated on FPGA.

### Accuracy Evaluation

The accuracy evaluation of the final mappings was performed using two methods; 1. reference software (*minimap2*) independent method that uses simulated reads where the exact mapping location is known; 2. reference software (*minimap2*) dependent method that considers real reads mapped using original *minimap2* to be the truth set.

#### Using simulated reads

For nanopore reads, the accuracy of *minimap2-fpga* is near-identical to the accuracy of original *minimap2* for both with base-level alignment (Figure 1e, Supplementary Table S1) and without base-level alignment (Figure 1f, Supplementary Table S1), on both computer systems (Intel FPGA and Xilinx FPGA systems), as demonstrated by the accuracy curves overlapping one another. In Figure 1e and Figure 1f, the horizontal axis shows the accumulative mapping error rate of mapped reads (accumulative number of wrong mappings / accumulative number of mapped reads) and the vertical axis shows the fraction of mapped reads (accumulative number of mapped reads / total number of mapped reads) corresponding to the error rate.

Similar to nanopore reads, the accuracy for PacBio reads from *minimap2-fpga* is also near-identical to that of original *minimap2* (Table 1). The mapping quality (MAPQ) score distribution for PacBio reads had only 3 distinct values. The top half of Table 1 contains the accuracy evaluation for sequence mapping with base-level alignment enabled and the bottom half is for sequence mapping without base-level alignment. In Table 1, the first column shows the mapping quality threshold (MAPQ_T). For each system in Table 1, the first column gives the number of reads mapped with MAPQ>=MAPQ_T and the second column gives the number of wrong mappings included in those mappings. For both with and without base-level alignment, the number of reads mapped with MAPQ>=MAPQ_T by *minimap2-fpga*, is equal to or even slightly higher than that of *minimap2*. The number of unmapped reads was the same on both *minimap2* and *minimap2-fpga*.

**Table 1.**
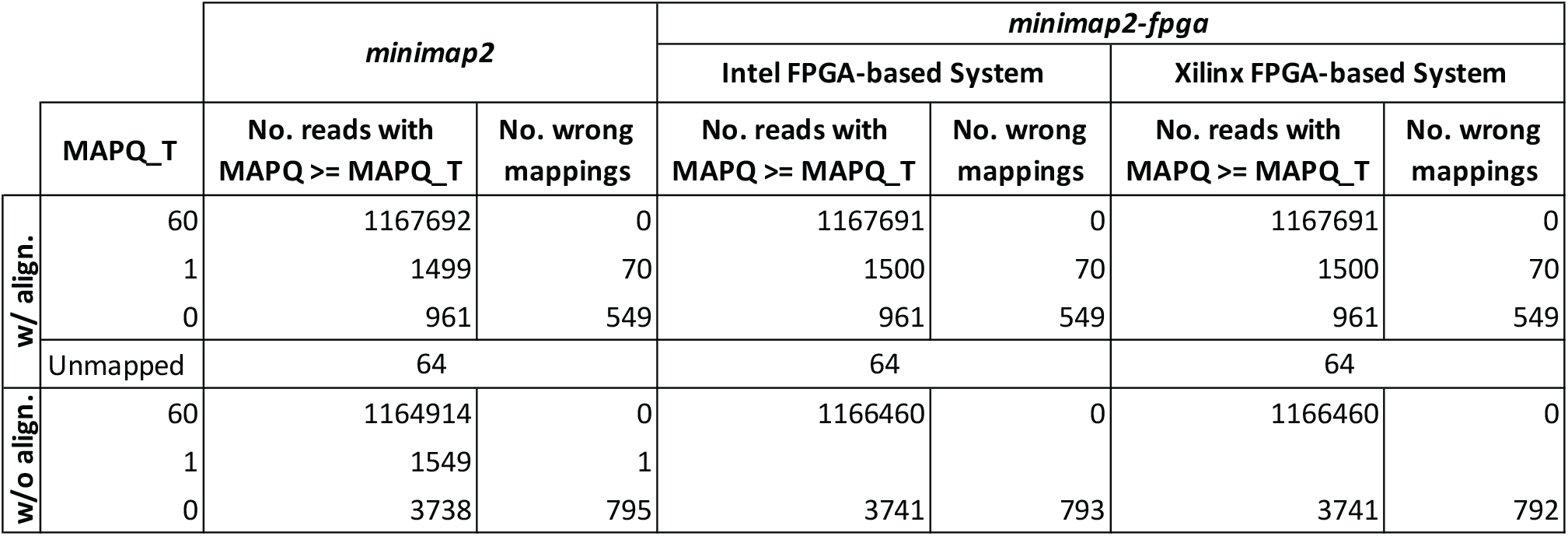
Accuracy comparison with PacBio simulated reads.

#### Using real reads

For the ∼ 30× ONT dataset, 23,228 out of 9,083,052 reads were unmapped in both versions of *minimap2* (*minimap2* and *minimap2-fpga*) on both Intel FPGA-based system and the Xilinx FPGA-based system (Table 2). On the Intel FPGA-based system, 9,045,287 mappings were the same in both versions of *minimap2*, making the output similarity between the two versions 99.58%. On the Xilinx FPGA-based system, 9,053,718 mappings were the same in both versions of *minimap2*, making the output similarity between the two versions 99.68%.

**Table 2.** Accuracy comparison with real reads.

|  | Intel FPGA-based System |  | Xilinx FPGA-based System |  |
| --- | --- | --- | --- | --- |
|  | ONT | PacBio | ONT | PacBio |
| **Total reads** | 9,083,052 | 11,714,594 | 9,083,052 | 11,714,594 |
| **Unmapped in both** | 23,228 | 62,461 | 23,228 | 62,492 |
| **Same** | 9,045,287 | 11,625,209 | 9,053,718 | 11,635,943 |
| **Mismatches** | 14,535 | 26,883 | 6,106 | 16,159 |
| **Only in *minimap2*** | 2 | 10 | 0 | 0 |
| **Only in *minimap2-fpga*** | 0 | 31 | 0 | 0 |
| **Same %** | 99.58 | 99.24 | 99.68 | 99.33 |

For the ∼ 50× PacBio dataset, 62,461 and 62,492 out of 11,714,594 reads were unmapped in both versions of *minimap2* on Intel FPGA-based system and the Xilinx FPGA-based system, respectively. On the Intel FPGA-based system, 11,625,209 mappings were the same in both versions of *minimap2*, making the output similarity between the two versions 99.24%. On the Xilinx FPGA-based system, 11,635,943 mappings were the same in both versions of *minimap2*, making the output similarity between the two versions 99.33%.

The output similarity between *minimap2* and *minimap2-fpga* is higher on the Xilinx FPGA-based system than on the Intel FPGA-based system due to the higher maximum loop trip count (H=1024 vs H=512) possible on Xilinx FPGA-based system (detailed in Materials and Methods).

## Discussion

### Previous Acceleration Work

Second-generation sequence analysis has been extensively optimized through various approaches, including CPU-based optimizations^9,10^, GPU-based optimizations^11^ and FPGA-based optimizations^12,13^ and custom MPSoC designs^14^. FPGA-based accelerations include a work^12^ that achieves a speedup over second-generation sequence analysis tool - Bowtie^15^ in both single-core and 16-core performance, a work^16^ that accelerates BWA-SW aligner^17^, and an end-to-end integrated tool^13^ with output accuracy comparable to BWA-MEM^18^. Commercial FPGA-based solutions, such as the DRAGEN system accompanying Illumina sequencers, have also emerged, benefiting from prior academic works in this field.

The emergence of third-generation sequencing, which offers massive throughput and ultra-long reads, has amplified the need for accelerated sequence analysis. CPU-based optimizations for third-generation sequence analysis include the acceleration of base-level alignment of *minimap2* on three processors^19^, multi-stage acceleration of *minimap2* on modern CPUs with Intel AVX-512 instructions^3^. GPU-based optimizations include the acceleration of adaptive banded event alignment in ONT raw signal analysis^20^ and acceleration of chaining step of *minimap2*^4^. FPGA-based solutions include acceleration of the base-calling task in Oxford Nanopore sequence analysis^21^, an integration^22^ of the GACT-X aligner architecture^23^ with *minimap2*, acceleration of *minimap2*’s chaining step^5,6^ and acceleration of selective genome sequencing^24^.

### Challenges in Hardware-Software Integration

Accelerating an isolated section of a large software tool on hardware accelerator platforms is commonly seen in the domain of hardware acceleration. However, integrating such accelerators with the rest of the software tools is less common and can pose significant challenges. A naive integration could diminish the astounding speed-up gained by the accelerator alone for the isolated portion of the software tool, and in some cases, even slow it down. Overcoming these challenges is necessary to achieve a speed-up in the end-to-end time. However, in the bioinformatics domain, integrating such accelerators with the software tool is crucial to provide a complete genomics analysis flow that can run with the accelerated tool.

*minimap2-fpga* is an end-to-end FPGA accelerated version for the gold-standard third-generation genomics sequence alignment tool *minimap2*. This integration resulted in a significant speed-up in the end-to-end time of the tool, while preserving the accuracy of the final output. Achieving this speed-up while maintaining accuracy required careful fine-tuning of all stages of the hardware-software processing pipeline, which involved addressing several engineering challenges. To enhance the accuracy of the final output while staying within the FPGA’s logic resource constraints, we made novel architectural changes to our previous hardware accelerator for isolated chaining^6^. We also created linear-regression based models to predict the time taken for each chaining task on hardware and software, allowing for more intelligent task-splitting decisions. To optimize access to a smaller number of hardware accelerator units from a larger number of software threads, we developed a scheduling mechanism.

### OpenCL for Accelerating Bioinformatics Kernels

We utilized OpenCL High-Level Synthesis (HLS) instead of a Hardware Descriptive Language (HDL) such as Verilog or VHDL for hardware accelerator implementation. HLS allowed for the accelerator to be designed using a high-level language similar to the C programming language, with the HLS compiler being considerably guided manually by appropriate compiler directives to generate the desired hardware. While using HDL instead of HLS could have improved the performance and FPGA resource usage of the accelerator by allowing for fine-tuning and optimization at a finer granularity, using HLS provided greater flexibility in making changes to an already designed hardware with significantly less development time. For instance, when the chaining step accelerator in this work needed to be modified from the previous version^6^ to the current version to improve the accuracy of the output it generates, the changes could be implemented (after the architectural changes were decided at a higher level) with relatively lesser time and effort. If this were to be done with HDL, it would require significantly more time and effort to implement, verify and debug the additional features added to the accelerator. This is particularly beneficial for bioinformatics software like *minimap2*, which is fast-evolving and requires quick implementation of future algorithm changes in the hardware.

### Future Work

Once the chaining step of *minimap2* is accelerated by ∼ 4× (on average) in this work, the other sections of the tool such as *ksw_extd2_sse41, rs_sort_128x* have now become the hotspots (see Figure 1d). To get even better improvements in the tool’s end-to-end time, future research can design hardware accelerators for these sections as well and integrate the accelerators into the rest of the software using the same techniques discussed in this work. In addition to creating specialized hardware, some work^3,25^ have already improved the software performance of core algorithms used in these remaining hotspots and such work can also be integrated into our work to get better performance improvements in end-to-end time. Furthermore, for sequence analysis applications such as selective sequencing, which sequences specific regions or targets of interest within the genome, *minimap2* is employed without base-level alignment (e.g. *readfish*^26^). *minimap2-fpga*, which offers higher speed-ups for mapping tasks that do not require base-level alignment, can be used as a replacement for *minimap2* to achieve speed-ups in such selective sequencing applications.

### Materials and Methods

### Computer Systems and Datasets

To evaluate the performance and accuracy with real sequencing data, two publicly available datasets were used: a NA12878 sample sequenced on an ONT PromethION sequencer (SRA Accession no. SRX11368472, 9.1M reads, 93.4 Gigabases^27^); and an HG002/NA24385 sample sequenced on a PacBIO Sequel II (15+20 kb CCS, SRA Accession no. PRJNA586863, 11.7M reads, 166.2 Gigabases^28^). To evaluate accuracy using simulated sequencing data: 1 million synthetic ONT long reads were simulated from hg38 human reference genome using *squigulator*^29^ (ONT raw signal data in BLOW5 format^30^) followed by *buttery-eel*^31^ (basecall signal data); and, ∼ 1.2 million synthetic PacBio CCS reads were simulated using *pbsim*^32^ from hg38 reference.

The on-premise Intel FPGA-based system used for performance and accuracy benchmarking was a heterogeneous computing system comprised of a 20-core Intel Xeon Gold 6148 CPU (2.40 GHz) with 754 GB of RAM and an Intel Programmable Acceleration Card (Intel Arria 10 GX 1150 FPGA) with 8 GB DDR4 onboard memory. The cloud Xilinx FPGA-based system was an Amazon EC2 F1 instance with 8 vCPUs, 122 GiB instance memory and a Xilinx UltraScale+ VU9P based FPGA device.

### Performance and Accuracy Evaluation

To compare the runtime performance of *minimap2* and *minimap2-fpga*, the ONT ∼ 30× and the PacBio ∼ 50× datasets were mapped against hg38 human reference genome using *minimap2-fpga* and *minimap2*, with and without base-level alignment. Both *minimap2* and *minimap2-fpga* were run with 8 CPU threads, on both cloud Xilinx FPGA system and on-premise Intel FPGA system (Supplementary Note 4).

To evaluate accuracy using simulated sequencing data, the generated simulated reads were mapped to the hg38 human reference genome using *minimap2-fpga* and *minimap2*, with and without base-level alignment. The outputs from both the computer systems were then evaluated using the *mapeval* utility in *paftools* scripts available under the *minimap2* repository. For real data (ONT ∼ 30× and the PacBio ∼ 50× datasets), the mapping output generated by original *minimap2* (with base-level alignment) software was considered the truth-set. The output generated by *minimap2-fpga* (with base-level alignment) for real data was compared against the truth-set using a SAM comparison utility^33^ (Supplementary Note 4).

### *minimap2-fpga* development, optimisation and integration

*minimap2-fpga* was implemented using C/C++ and OpenCL. To accelerate the compute-intensive chaining step, *minimap2-fpga* utilizes either the custom FPGA-based hardware accelerator (implemented using OpenCL HLS, extending our previous work^6^) or the CPU-based software implementation, based on the predicted hardware/software execution times of each chaining task. To perform these predictions, a linear regression model was trained by analyzing the hardware/software execution times of a set of chaining tasks measured with a representative dataset. A shell script has been provided under the *minimap2-fpga* repository to execute this one-time linear regression model training process on a different dataset if deemed necessary in future.

Intel Arria 10 GX FPGA Acceleration Stack 1.2.1 (with Intel FPGA SDK for OpenCL 19.4 and Intel Quartus Prime Pro 19.2) was used for hardware accelerator development on the on-premise Intel FPGA-based system (Supplementary Note 1). On the cloud Xilinx FPGA system, AWS EC2 FPGA Development Kit with Xilinx Vitis v2021.2 was used for the hardware accelerator development. For Xilinx FPGA, the OpenCL HLS pragmas used in the Intel FPGA hardware accelerator implementation had to be converted to supported Xilinx OpenCL HLS pragmas. Due to the lack of a pragma in Xilinx OpenCL HLS for data prefetching from global memory to local memory (line 7-8 in Supplementary Algorithm 1), this prefetching was manually implemented.

We enhanced our previous hardware accelerator^6^, to achieve a level of accuracy in the final mapping output that is comparable to that from the original *minimap2* software. When we naively integrated our previous hardware accelerator^6^ to *minimap2*, the end-to-end accuracy was considerably lower in comparison to the original *minimap2* (Supplementary Figure S1). Further investigation revealed that setting a maximum loop trip count in the chaining algorithm to 64 was the culprit (H=64 in lines 18-24 of Supplementary Algorithm 2). To improve the accuracy, the maximum trip count had to be increased. However, naively increasing this trip count considerably increased the resource usage on the FPGA which resulted in a design that could not fit the available FPGA area (in terms of the number of logic components such as LUTs, FFs and DSPs). To improve accuracy while keeping the resource usage low, we restructured the affected inner loop into subpartitions (detailed in Supplementary Note 1). With this improvement to the hardware accelerator, a maximum trip count of 512 (i.e. H=512) and 1024 (i.e. H=1024) were achieved on the Intel FPGA and the Xilinx FPGA accelerator, respectively. A single hardware kernel (i.e. N=1 in Supplementary Figure S2) was configured on both the Intel and Xilinx FPGA accelerators. The overall accuracy is now near-identical to the original *minimap2* (Figure 1e and 1f).

To achieve further speed up (∼ 4× on average) in the chaining step while retaining the output accuracy, further optimisations were performed. To process chaining tasks on the FPGA-based hardware accelerator, the input chaining anchor data and the output chaining scores need to be transferred to/from the FPGA device connected to the host CPU via PCI Express interface. However, this data transfer incurs overhead, which is only justifiable if the execution time on the hardware accelerator is significantly smaller than the software execution time. Therefore, chaining tasks are processed on the FPGA-based hardware accelerator only if the total processing time (data transfer time + execution time) on hardware is smaller than the execution time on software. In this work, the hardware-software split of chaining tasks presented in our previous work^6^ was optimised with a novel approach that is based on predicted hardware/software processing times. Equation 1 and 2 depict the linear regression models formulated to predict the processing time of a given chaining task on hardware and software respectively. In these equations, *n* is the number of anchors and *K*_1_, *K*_2_, *C*_1_, *C*_2_ are constants. More details on how the equations were derived are in Supplementary Note 2.

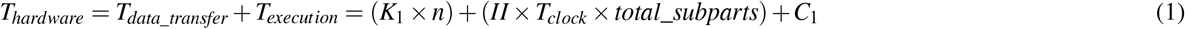

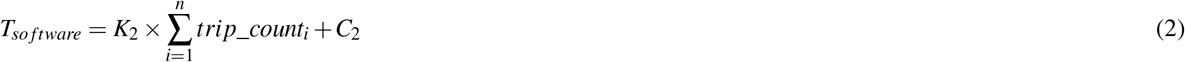

Based on the hardware/software execution time predictions, an improved scheduling algorithm was created to schedule chaining tasks on hardware, so that the waiting time to access the smaller number of hardware kernels (configured on the FPGA with limited logic resources) from a larger number of software threads (available in a typical a high-performance computing system) is minimized. More details on hardware-software integration and hardware scheduling are in Supplementary Note 3.

## Conclusion

In this work, we presented *minimap2-fpga*, a hardware-accelerated version of *minimap2* that integrates an FPGA kernel optimized for chaining. *minimap2-fpga* supports mapping DNA reads from both ONT and PacBio onto a reference genome. For end-to-end execution, *minimap2-fpga* was up to 79% and 53% faster than original *minimap2* for ONT and PacBio datasets respectively, when mapping without base-level alignment. When mapping with base-level alignment, *minimap2-fpga* was up to 62% and 10% faster than *minimap2* for ONT and PacBio datasets respectively. The mapping output accuracy of *minimap2-fpga* was near-identical to that of *minimap2*. To achieve this, we had to address scheduling overheads in multi-threaded execution and resolve an unexpected issue that affected mapping accuracy, which only became apparent during full integration. Our work highlights the importance of integrating accelerated kernels back into software workflows and offers valuable insights for other bioinformatics tools requiring heterogeneous computing accelerations.

## Data Availability

The experimental data used in benchmarking experiments are from publicly available datasets on NCBI Sequence Read Archive (ONT data: SRX11368472, PacBio data: PRJNA586863). Simulated reads can be created by following the instructions in Supplementary Note 4.2.

## Acknowledgements

We thank D. Lin and W. Kaplan for providing exceptional technical assistance to leverage the institution’s powerful computing resources, particularly the Intel FPGA-equipped systems, in unique and innovative ways. K.L. is supported by Australian Government Research Training Program (RTP) Scholarship. H.G. is supported by Australian Research Council DECRA Fellowship DE230100178. This research was supported (partially) by the Australian Government through the Australian Research Council’s Discovery Projects funding scheme (project DP230100651). The views expressed herein are those of the authors and are not necessarily those of the Australian Government or the Australian Research Council.

## Author contributions statement

K.L. conducted the hardware/software development/optimisations and experiments under the supervision of H.G. and S.P. H.S. contributed to modifying the hardware to enhance accuracy and collaborated on software development. K.L. and H.G. wrote the manuscript with input from H.S. and S.P.

## Additional information

H.G., has previously received travel and accommodation expenses to speak at Oxford Nanopore Technologies conferences.

H.G. has had paid consultant roles with Sequin PTY LTD. The authors declare no other competing financial or non-financial interests.

