## supplementary_material for "*minimap2-fpga*: Integrating hardware-accelerated chaining for efficient end-to-end long-read sequence mapping"

#### List of Tables

S1 Accuracy comparison with ONT simulated reads . . . . . 2

#### List of Figures

S1 Accuracy of *minimap2-fpga* when combined with previous hardware accelerator[6] on the Intel FPGA-based system 3
S2 The architecture of the FPGA-based hardware accelerator designed for *minimap2*'s chaining step acceleration . . 4

#### Supplementary Notes

**1 FPGA-based Chaining Step Accelerator 4**
**2 Hardware-Software Split 6**
**3 Hardware-Software Integration 8**
**4 Experiments and Benchmarks 10**
4.1 Performance Benchmarking . . . . . 10
4.2 Accuracy Benchmarking . . . . . 10

**Table S1.** **Accuracy comparison with ONT simulated reads**

| | | *minimap2* | | *minimap2-fpga* | | | |
|---|---|---|---|---|---|---|---|
| | | | | Intel FPGA-based System | | Xilinx FPGA-based System | |
| | MAPQ_T | No. reads with MAPQ >= MAPQ_T | No. wrong mappings | No. reads with MAPQ >= MAPQ_T | No. wrong mappings | No. reads with MAPQ >= MAPQ_T | No. wrong mappings |
| w/ base-level alignment | 60 | 960462 | 0 | 960470 | 0 | 960464 | 0 |
| | 59 | 10210 | 1 | 10213 | 1 | 10209 | 1 |
| | 38 | 20705 | 1 | 20694 | 1 | 20704 | 1 |
| | 8 | 1477 | 1 | 1477 | 1 | 1477 | 1 |
| | 5 | 492 | 2 | 492 | 2 | 492 | 2 |
| | 4 | 492 | 1 | 492 | 1 | 492 | 1 |
| | 3 | 546 | 5 | 544 | 5 | 544 | 5 |
| | 2 | 568 | 2 | 570 | 2 | 570 | 2 |
| | 1 | 2730 | 456 | 2730 | 456 | 2730 | 456 |
| | 0 | 2198 | 1241 | 2198 | 1241 | 2198 | 1241 |
| | Unmapped | 120 | | 120 | | 120 | |
| w/o base-level alignment | 60 | 970514 | 0 | 970515 | 0 | 970516 | 0 |
| | 22 | 16981 | 1 | 16981 | 1 | 16979 | 1 |
| | 6 | 1193 | 1 | 1194 | 1 | 1195 | 1 |
| | 5 | 1176 | 3 | 1177 | 3 | 1177 | 3 |
| | 4 | 1107 | 1 | 1106 | 1 | 1104 | 1 |
| | 3 | 1131 | 9 | 1130 | 9 | 1130 | 8 |
| | 2 | 1223 | 10 | 1222 | 10 | 1222 | 10 |
| | 1 | 1525 | 22 | 1525 | 22 | 1525 | 22 |
| | 0 | 5030 | 1985 | 5030 | 1985 | 5032 | 1985 |

The top half is for SAM outputs generated with base-level alignment enabled, and the bottom half is for PAF outputs generated without base-level alignment. The first column is mapping quality threshold (MAPQ_T). For each system, the first column shows the number of reads mapped with MAPQ>=MAPQ_T and the second column shows the number of wrong mappings in those reads.

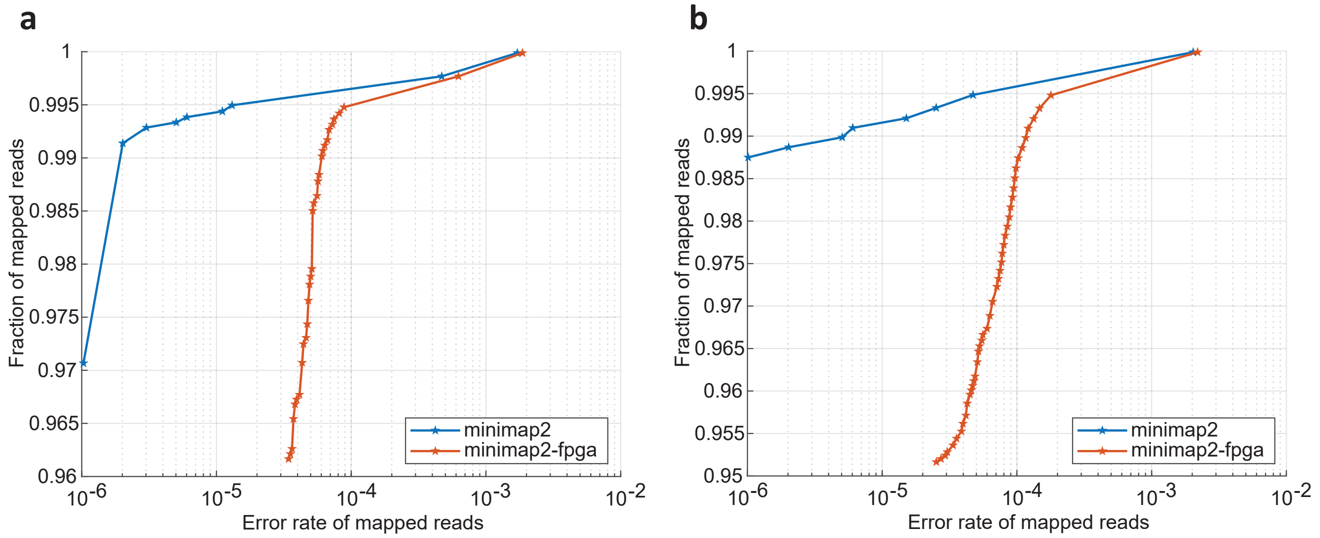


**Figure S1.** **Accuracy of *minimap2-fpga* when combined with previous hardware accelerator [6] on the Intel FPGA-based system**

(**a**) Accuracy comparison performed with simulated ONT reads for original *minimap2* vs. *minimap2-fpga* with base-level alignment. (**b**) Accuracy comparison performed with simulated ONT reads for original *minimap2* vs. *minimap2-fpga* without base-level alignment.

### 1 FPGA-based Chaining Step Accelerator

The FPGA-based hardware accelerator architecture in our previous work [6] was extended as shown in Figure S2. With the modifications to the hardware accelerator, it was able to achieve better accuracy (compare Supplementary Figure S1 vs. Figure 1e, 1f) in the chaining output while keeping the FPGA resource usage low (so that the accelerator could fit in the limited FPGA area available to use). Furthermore, we successfully maintained the desired speed-up factor throughout the optimization process. Algorithm 1 describes the HLS algorithm used to generate the modified hardware architecture in Figure S2.

*minimap2*'s software chaining algorithm (Algorithm 2) computes $H$ (at most) chaining scores for each anchor, using previous $H$ (at most) anchors (*for* loop in lines 17-23 of Algorithm 2). In our previous hardware accelerator implementation [6], we parallelized this computation by utilizing $H$ score computation units to perform the computations in parallel. Considering the FPGA resource limitations and *minimap2*'s original algorithm definitions, $H$ was set to 64. However, to achieve a mapping accuracy similar to software implementation of *minimap2*, $H$ needed to be increased. Increasing the number of parallel score computation units ($H$) directly to achieve better accuracy, resulted in a design that couldn't fit in the FPGA being used. To resolve this resource utilization issue, the hardware architecture is modified so that the $H$ score computations corresponding to a single anchor are performed in $M$ sub-parts (see Figure S2 and Algorithm 1). Score computations in a sub-part are performed in parallel with only $P$ score computation units (note that previous accelerator [6] needed $H$ score computation units). As most recently used $H$ anchors ($A[i]$) and the maximum score values corresponding to them ($F[i]$), are needed for subsequent score computations, $H = M \times P$ anchors and their maximum score values are stored in first-in, first-out (FIFO) buffers implemented with hardware shift registers.

The modified hardware accelerator with additional control logic was optimized to have an Initiation Interval (II) of 3, making it able to process a sub-part every 3 clock cycles.

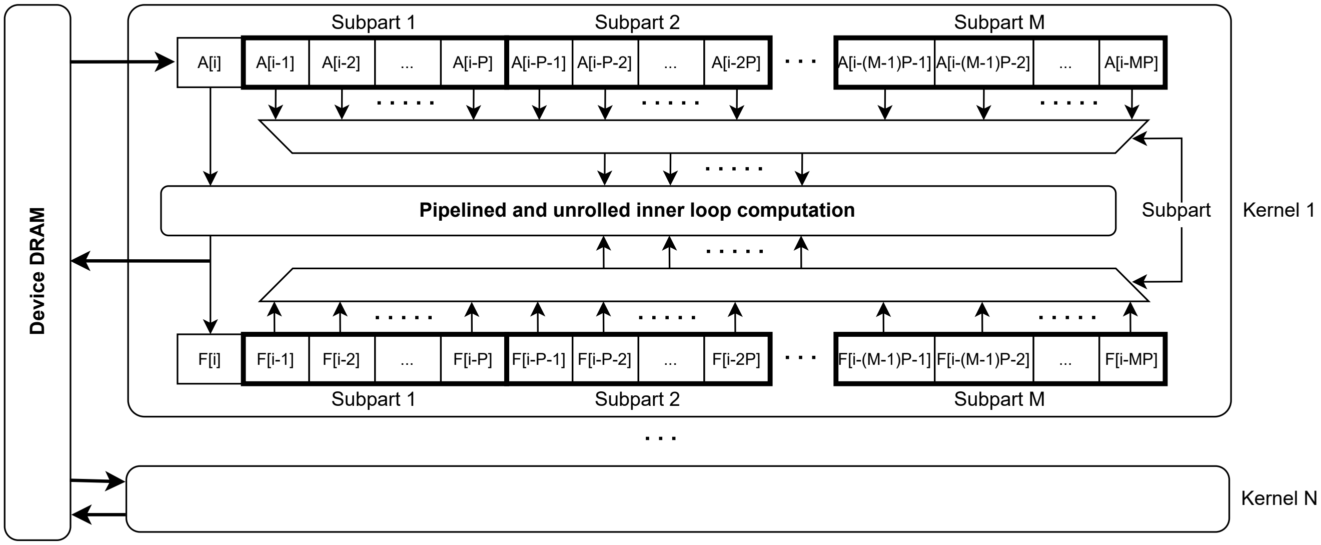


**Figure S2. The architecture of the FPGA-based hardware accelerator designed for *minimap2*'s chaining step acceleration**
The hardware architecture of our previous hardware accelerator [6] was modified to achieve better accuracy while preserving the speed-up when compared to original *minimap2* software.

**Algorithm 1** Hardware Accelerator's Behavior (HLS)

```
Inputs: a[], num_subparts[], total_subparts, q_span, avg_qspan_scaled, max_dist_x, max_dist_y, bw
Outputs: f[], p[]
A[M * P + 1] = {0}, F[M * P + 1] = {0}
i = 0
subparts_processed = 0
subparts_to_process = 0
for g = 0; g < total_subparts; g++ do
    if subparts_processed == 0 then
        A[0] = __prefetching_load(&a[i])
        subparts_to_process = __prefetching_load(&num_subparts[i])
    end if
    max_f = q_span
    max_j = -1
    #pragma unroll
    for j = P; j > 0; j-- do
        buffer_offset = subparts_processed * P
        score = compute_score(A[i], A[buffer_offset + j], F[buffer_offset + j], max_dist_x, max_dist_y, bw, q_span,
                    avg_qspan_scaled)
        if score >= max_f and score != q_span then
            max_f = score
            max_j = i - j - buffer_offset
        end if
    end for
    if max_f > F[0] then
        F[0] = max_f
        f[i] = max_f
        p[i] = max_j
    end if
    subparts_processed++
    #pragma unroll
    for reg = M * P; reg > 0; reg-- do
        if subparts_processed == subparts_to_process then
            A[reg] = A[reg - 1]
            F[reg] = F[reg - 1]
        end if
    end for
    if subparts_processed == subparts_to_process then
        i++
        F[0] = 0
        subparts_processed = 0
    end if
end for
```

#### 2 Hardware-Software Split

A novel hardware-software split mechanism was introduced to process the chaining tasks either on hardware accelerator or the CPU (as software) based on their computation times predicted with theoretical models. Equation 1 and Equation 2 give the models derived to predict the time taken for a given chaining task to run on hardware accelerator and on CPU (as software) respectively.

The time taken to process a given chaining task on hardware ($T_{hardware}$, given in Equation 1) is the sum of the times taken for data transfer ($T_{data\_transfer}$ = input transfer from host to the hardware accelerator + output results transfer from hardware accelerator to the host) and the execution time on the hardware accelerator ($T_{execution}$). The data transfer time ($T_{data\_transfer}$) is proportional to the number of anchors being transferred ($n$). As the output for a single subpart is generated by the accelerator every $II$ clock cycles, the execution time in terms of clock cycles is $II \times total\_subparts$. Multiplying the clock cycles by the clock period ($T_{clock}$) gives the execution time in seconds ($T_{execution}$). The constant $C_1$ accounts for the initial pipeline delay and the overhead time taken for OpenCL API calls by the host.

To derive the relationship between the time taken to process a given chaining task on the CPU as software ($T_{software}$, as given in Equation 2), it is necessary to refer to the original software chaining algorithm. This algorithm is presented in Algorithm 2 (also presented in our previous work[6]). The number of computational steps can be estimated by the total number of iterations (i.e. total trip count) taken by the *for* loop in Lines 17-23 of Algorithm 2 and is represented by $\sum_{i=1}^{n} trip\_count_i$ in Equation 2. Computing the total trip count corresponding to a single anchor can be done prior to the real execution of the *for* loop by using the *while* loop in Lines 13-15 of Algorithm 2 (this loop computes the starting index ($st$) used in the *for* loop in Lines 17-23 and $min(i-H, st)$ gives the trip count corresponding to the anchor). Summing up the trip count values corresponding to all the anchors gives $\sum_{i=1}^{n} trip\_count_i$ in Equation 2. $C_2$ in Equation 2 accounts for the overhead setup time for the *for* loop in Lines 17-23 of Algorithm 2.

Based on a one-time experiment done with a representative dataset, the values for the coefficients in Equation 1 and 2 (i.e. $K_1$, $K_2$, $C_1$, $C_2$) are found. After, $T_{hardware}$ and $T_{software}$ are calculated for each chaining task at the run time. If $T_{hardware} < T_{software}$, the chaining task is decided to be processed on hardware (see Supplementary Note 3 for hardware scheduling details), otherwise, the chaining task is processed on the host CPU itself as software.

**Algorithm 2** *minimap2*'s Chaining Algorithm

```
Inputs: a[], n, max_dist_x, max_dist_y, bw
Outputs: f[], p[]
▷ Calculate average seed length (avg_qspan)
sum_qspan = 0
for i = 0; i < n; i++ do
    sum_qspan += a[i].y >> 32 & 0xff
end for
avg_qspan = sum_qspan / n
st = 0                                                  ▷ Starting index of inner loop
for i = 0; i < n; i++ do
    ▷ Initialize best score (max_f) and predecessor (max_j)
    max_f = q_span = a[i].y >> 32 & 0xff
    max_j = -1
    ▷ Find the starting index (st) of inner loop
    while st < i and a[i].x > a[st].x + max_dist_x do
        st++
    end while
    ▷ Compute scores between anchor i and previous H (at most) anchors and find the maximum score (max_f)
    for j = i - 1; j >= st and j >= i - H; j-- do
        score = compute_score(a[i], a[j], f[j], avg_qspan,
                              max_dist_x, max_dist_y, bw)
        if score > max_f then
            max_f = score
            max_j = j
        end if
    end for
    ▷ Store maximum score and best predecessor for anchor i
    f[i] = max_f
    p[i] = max_j
end for
```

#### 3 Hardware-Software Integration

The FPGA-based hardware accelerator is carefully integrated to the *minimap2* original software while minimally affecting the multi-threaded behavior of original software so that an efficient reduction in tool's end-to-end runtime can be achieved. OpenCL API functions are used for data transfer and communication between the host CPU (software) and the FPGA-based accelerator (hardware).

*minimap2* processes DNA reads on software with multiple threads by using a fork-join thread model. This allows *minimap2* to parallelize the computations being performed while maximally utilizing the multiple cores available in the CPU. When a chaining request is issued by any of the software threads of hardware-software integrated version of *minimap2* (i.e. *mm_chain_dp* function is called by a software thread), it is decided whether the requested chaining task should be processed on hardware or software with the hardware-software split mechanism discussed in Supplementary Note 2. If the chaining task is decided to be processed on software, the associated software thread continues to execute the chaining task on the CPU itself.

Since the FPGA devices are resource-constrained, the number of hardware kernels (*N* in Fig. S2) that can be configured on the device is usually lower than the number of software threads available on a typical high performance computing (HPC) system. Therefore, when the chaining task is decided to be processed on hardware, each software thread in the host CPU tries to schedule the chaining computation on one of the *N* hardware kernels configured on the FPGA device as given in Algorithm 3.

In lines 1-18 of Algorithm 3, given the predicted hardware and software execution times, the algorithm tries to find a hardware kernel on which the total time to finish processing the chaining task (*total_hw_time_pred*, which is the sum of the predicted hardware processing time (*hw_time_pred*) and the waiting time to access that kernel (*wait_time*)), is smaller than the predicted software execution time (*sw_time_pred*). If *total_hw_time_pred* is greater than or equal to *sw_time_pred*, the associated software thread executes the chaining task on the CPU itself. If *total_hw_time_pred* is smaller than *sw_time_pred*, the hardware kernel which satisfied the condition is recorded (*kernel_id*) and the chaining task is inserted into a first-in, first-out (FIFO) queue corresponding to the hardware kernel (*hw_queue[kernel_id]*), so that the CPU can wait and execute the task on the hardware kernel when the chaining task comes first in the queue. Lines 20-33 of Algorithm 3 correspond to the section where it takes the chaining task out from the queue and processes it on the previously recorded hardware kernel (*kernel_id*) when it is ready to do so. As this algorithm is performed on multiple software threads running in parallel and the *N* hardware kernels and the *N* hardware queues are shared among the multiple threads, necessary synchronization mechanisms (highlighted in blue color in Algorithm 3) are implemented with two mutex locks for each hardware kernel - one for accessing/modifying the queue corresponding to the kernel (called "queue lock") and one for accessing the kernel for chaining task execution (called "execution lock").

**Algorithm 3** Hardware Scheduling Algorithm

```
Inputs: hw_time_pred, sw_time_pred
should_wait = false
for kernel_id = 0; kernel_id < NUM_HW_KERNELS; kernel_id++ do
    Acquire kernel's queue lock
    wait_time = hw_end_times[kernel_id] - current_time
    total_hw_time_pred = wait_time + hw_time_pred
    if total_hw_time_pred < sw_time_pred then
        hw_queue[kernel_id].push(tid)
        end_times[kernel_id] = current_time + total_hw_time_pred
        should_wait = true
    end if
    Release kernel's queue lock
    if should_wait == true then
        break
    end if
end for
if should_wait == false then
    "process the chaining task on software"
end if

got_in = false
while true do
    Acquire kernel's execution lock
    Acquire kernel's queue lock
    if hw_queue[kernel_id].front() == tid then
        hw_queue[kernel_id].pop()
        got_in = true
    end if
    Release kernel's queue lock
    if got_in == true then
        break
    end if
    Release kernel's execution lock
end while
"process the chaining task on hardware"
Release kernel's execution lock
```

### 4 Experiments and Benchmarks

#### 4.1 Performance Benchmarking

The commands used for the performance benchmarking detailed under "Performance Comparison" are given below. The same commands were used both on Intel FPGA-based system and the Xilinx FPGA-based system.

For performance benchmarking with ONT data (w/o alignment and w/ alignment respectively):

```
$ minimap2_fpga -x map-ont -t 8 hg38noAlt.idx ont_reads.fastq > alignment.paf
$ minimap2_fpga -ax map-ont -t 8 hg38noAlt.idx ont_reads.fastq > alignment.sam
```

For performance benchmarking with PacBio CCS data (w/o alignment and w/ alignment respectively):

```
$ minimap2_fpga -x asm20 -t 8 hg38noAlt.idx pbccs_reads.fastq > alignment.paf
$ minimap2_fpga -ax asm20 -t 8 hg38noAlt.idx pbccs_reads.fastq > alignment.sam
```

#### 4.2 Accuracy Benchmarking

The commands used for generating the simulated reads used in "Accuracy Evaluation" are given below.

For ONT simulated reads generation:

```
# generate raw signal data
$ squigulator hg38noAlt.fa -x dna-r9-prom -o ont_sim_reads.blow5 -n 1000000 -t 8
# basecall the raw signal data
$ buttery-eel  -i  ont_sim_reads.blow5  --guppy_bin /path/to/ont-guppy-6.3.7/bin/
 --config dna_r9.4.1_450bps_hac_prom.cfg -x cuda:all -o ont_sim_reads.fastq
 --port 5555 --use_tcp
```

For PacBio simulated reads generation:

```
# generate reads
$ pbsim --data-type CCS --length-max 20000 --depth 6 --model_qc data/model_qc_ccs
--length-mean 15000 hg38noAlt.fa
# convert reads to a mapeval compatible format
$ k8-Linux paftools.js pbsim2fq hg38noAlt.fa.fai <reads>.maf > pbccs_sim_reads.fa
```
